# Reactive Oxygen Species Regulate the Inflammatory Function of NKT Cells through Promyelocytic Leukemia Zinc Finger

**DOI:** 10.1101/122390

**Authors:** Yeung-Hyen Kim, Ajay Kumar, Cheong-Hee Chang, Kalyani Pyaram

## Abstract

Reactive oxygen species (ROS) are byproducts of aerobic metabolism and contribute to both physiological and pathological conditions as second messengers. ROS are essential for antigen specific activation of T cells, but little is known about what role ROS play in NKT cells. In the current study, we investigated the role of ROS in NKT cell function. We found that ROS levels are similar among CD4, CD8 and NKT cell subsets in the thymus. However, NKT cells, but neither CD4 nor CD8 T cells, showed dramatically increased ROS in the spleen and liver but not in adipose tissues. ROS in the peripheral NKT cells were primarily produced by NADPH oxidases not mitochondria. Accordingly, ROS-high NKT cells were susceptible to oxidative stress and underwent apoptotic cell death. Furthermore, ROS play an important role in regulating the inflammatory function of NKT cells because antioxidant treatment of NKT cells showed reduced frequencies of IFN-γ^+^ and IL-17^+^ cells. In line with this, freshly isolated ROS-high NKT cells had more NKT1 and NKT17 cells but less NKT2 than ROS-low cells. These characteristics are regulated by promyelocytic leukemia zinc finger (PLZF) as evidenced by low ROS in NKT cells from PLZF haplodeficient mice and also from adipose tissues that do not express PLZF. Conversely, ROS were highly elevated in CD4 T cells from mice ectopically expressing PLZF. Together, our study revealed for the first time that ROS regulate NKT cell functions through PLZF.

## Introduction

Invariant Natural Killer T (NKT) cells express highly restricted T cell receptor repertoire and share characteristics of both T cells and natural killer (NK) cells (1). NKT cells arise from a common precursor of CD4^+^CD8^+^ double positive thymocytes which have undergone TCR gene rearrangement and expression. NKT cells undergo progressive maturation after positive selection in the thymus. Using the expression pattern of CD24, NK1.1 and CD44, stage 0 (CD24^+^NK1.1^-^ CD44^-^), stage 1 (CD24^-^NK1.1^-^CD44^-^), stage 2 (CD24^-^NK1.1^-^CD44^+^) and stage 3 (CD24^-^NK1.1^+^CD44^+^) NKT cells have been identified (2). In addition, NKT cell functional subsets, NKT1, NKT2 and NKT17, were defined based on their distinct expression of transcription factors T-bet, GATA3 and RORγt together with promyelocytic leukemia zinc finger (PLZF) protein, respectively, (3). Thymus-derived NKT cells undergo further differentiation and functional specialization in the periphery and produce a broad range of cytokines and exhibit both pro-inflammatory and immunoregulatory characteristics (4).

NKT cells express PLZF that is a master transcription factor determining the innate T cell fate (5, 6). PLZF encoded by *Zbtb16* plays an important role in the acquisition of effector functions such as cytokine production and migration properties in NKT cells during developmental process (5, 6). PLZF is required for NKT cell development, and ectopic expression of PLZF in conventional T cells showed a phenotype and function similar to NKT cells supporting the critical role of PLZF (5-7). In addition, PLZF works as a negative regulator of cell cycle progression ultimately leading to growth suppression (8), and enforced expression of PLZF in myeloid cell lines resulted in inhibition of proliferation and differentiation (9). A recent study showed that PLZF promotes hepatic gluconeogenesis (10), suggesting a plausible role of PLZF in cell metabolism.

Unlike conventional T cells that undergo antigen dependent differentiation programming to become effector cells in the periphery, each effector NKT cell type is generated as they develop in the thymus (11). The signaling pathways that control cellular metabolism have been shown to have a crucial role in dictating the outcome of T cell activation and effector function (12). Distinct T cell subsets adopt specific metabolic programs to support their needs. Studies have suggested that changes in the metabolic profile of T cells are responsible for defining the effector functions and T cell subsets (13, 14). It remains to be investigated if NKT cell functions are also controlled by cell metabolism during their development in the thymus.

NKT cells play a unique and important role in many immune diseases including metabolic diseases such as autoimmune hepatitis (15-17) and obesity (18-20). It seems that the effector function of NKT cells is different depending on the tissue type. Hepatic NKT cells are known to induce inflammation, whereas NKT cells in adipose tissues possess a regulatory function that might have developed differently from hepatic NKT cells in the thymus (21). Currently, the role of cell metabolism controlling NKT cell functions is not well understood although studies have shown that disruption of Vps34 (a class III PI3 kinase) (22), metabolic regulator Fnip1 (23), or the autophagy pathway (24, 25) causes a defect in NKT cell development and function. NKT cells are also sensitive to a change in signaling pathways mediated by CD28 (26), ICOS (27), PI3K (28), PDK1, Wnt/β-Catenin (29) and mTOR (30, 31) all of which regulate the metabolic pathways.

Reactive oxygen species (ROS) are byproducts of aerobic metabolism which include superoxide anion (O_2_^-^), hydrogen peroxide (H_2_O_2_), and hydroxyl radicals (OH) (32). ROS mediate redox biology and oxidative stress, and contribute to both physiological and pathological conditions (32). ROS are essential second messengers in innate and adaptive immune cells (33). However, increased levels of ROS within immune cells can result in hyperactivation of inflammatory responses, resulting in tissue damage and pathology (32, 34). In the immune cells, ROS are known for killing the pathogens in the phagosomes through oxidative burst mediated by NADPH oxidase (NOX) (35). In T cells, ROS levels increase upon activation and the major initial source of ROS required for T cell activation is mitochondria (36). In the absence of ROS produced by mitochondria, antigen specific responses of T cells are compromised although they can proliferate under the lymphopenic environment (36). However, NOX can be invoked in response to mitochondria produced ROS to further sustain ROS levels so as to maintain T cell activation (37).

In the current study, we investigated the role of ROS in NKT cell functions. We found that ROS levels in NKT cells are dramatically increased compared to CD4 and CD8 T cells in the spleen and liver but not in the adipose tissues. As a consequence, NKT cells are susceptible to oxidative stress but not ER stress. Furthermore, the inflammatory function of NKT cells correlates with the levels of ROS evidenced by more NKT1 and NKT17 than NKT2 subsets among NKT cells with high ROS. Our study also revealed that ROS levels and ROS-mediated changes of NKT cells are controlled by PLZF.

## Materials and Methods

### Mice

PLZF transgenic mice driven by the *cd4* promoter (6), the *lck* promoter (7), PLZF deficient mice (38), and Vα14 TCR transgenic mice (39) have been previously described. All the mice were bred and maintained under specific pathogen-free conditions at the University of Michigan animal facility and used at 8-12 weeks of age. All animal experiments were performed under protocols approved by the University of Michigan Institutional Animal Care and Use Committee.

### Cell preparation and purification

Whole spleen and liver cells were mechanically disrupted onto 100 μm cell strainer to collect single cell suspension. Homogenized spleen cells were subjected to ammonium chloride potassium (ACK) lysis to remove red blood cells and washed, and then resuspended in PBS supplemented with 2% FBS (FACS buffer). Homogenized liver cells were resuspended in 40% isotonic Percoll solution, loaded on the top of a 70% Percoll solution (GE Healthcare), and then centrifuged at 970×g at room temperatures with no brakes for 30 minutes. Non-parenchymal cells were collected at the interface of the two Percoll layers, washed, and resuspended in FACS buffer. To sort NKT and CD4 T cells, spleen cells were incubated with synthetic peptide PBS57-loaded CD1d tetramers, anti-TCR-⁏ antibody and anti-CD4 antibody for 30 minutes on ice. NKT and CD4 T cells were then sorted using FACS Aria-III (BD). To sort ROS-low and ROS-high NKT cells, splenocytes were stained with DCFDA for 30 minutes prior to surface staining. Cells were sorted using FACS Aria-III (BD) to isolated ROS-high and ROS-low cells.

### Flow cytometry assay

The following antibodies were used: anti-mouse TCR-⁏ (H57-597) APC or Pacific Blue, PBS57 loaded CD1d tetramer APC or Pacific Blue, anti-mouse CD4 (GK1.5) PerCp-Cy5.5, anti-mouse CD8 (53-6.7) Am-cyan, anti-mouse NK1.1 (PK-136) PE-Cy7, anti-mouse CD44 (IM7) PerCp-Cy5.5, anti-mouse CD69 (H1-2F3) PE-Cy7, anti-mouse CD62L (MEL-14) eVolve605, anti-mouse IFN-γ (XMG1.2) FITC or APC, anti-mouse IL-4 (11B11) PE-Cy7, anti-mouse IL-17 (TC11-18H10) APC-eFluor780, anti-T-bet (eBio4B10) FITC, anti-RORγt (AFKJS-9) Pacific Blue, anti-GATA3 (L50-823) APC, and anti-PLZF (Mags-21F7) PE (all from eBioscience). Dead cells were excluded by staining with 1 μg/ml propidium iodide (Sigma-Aldrich). To measure intracellular cytokines, cells were stimulated for 5 hours with phorbol 12-myristate 13-acetate (PMA) (50 ng/ml, Sigma-Aldrich) and ionomycin (1.5 μM, Sigma-Aldrich) in the presence of Monensin (3 μM, Sigma-Aldrich), permeabilized using Cytofix/Cytoperm Plus (BD), then stained with the appropriate antibodies. Transcription factor staining to identify committed cells was performed using the Foxp3/transcription factor staining kit (eBioscience) and intranuclear staining for T-bet, RORγt, GATA3 and PLZF. Data were acquired on a FACS Canto II (BD) and analyzed using FlowJo (TreeStar software ver. 9.9).

### ROS detection

To measure total ROS, cells (1×10^6^/ml FACS tube) were incubated in RPMI media containing 10% fetal bovine serum (FBS) with 1 μM DCFDA (Invitrogen) for 30 minutes at 37°C. Cells were washed with FACS buffer and stained with relevant surface antibodies for flow cytometry. Dead cells were eliminated by staining with 1 μg/ml propidium iodide (Sigma-Aldrich). To measure ROS produced by mitochondria, 2.5 μM MitoSox (Invitrogen) was used. Data were acquired on a FACS Canto II (BD) and analyzed using FlowJo (TreeStar software ver. 9.9).

### Apoptosis assay

1×10^6^ total splenocytes and hepatocytes were resuspended in FACS buffer and stained for surface markers. Cells were analyzed for apoptosis after Annexin V staining according to the manufacturer’s instructions (BD).

### T cell activation

Sorted NKT cells were resuspended at 3×10^5^ cells per 96-well flat tissue culture plates in RPMI medium supplemented with 10% FBS with soluble α-Galcer (100 ng/ml) for 3 days on 37°C with 5% CO_2._ Sorted CD4 T cells (3×10^5^ cells/well) were stimulated with plate bound anti-CD3 antibody (5 μg/ml) with soluble anti-CD28 antibody (2 μg/ml). To measure intracellular cytokines expression, the cells were restimulated with PMA (50 ng/ml, Sigma-Aldrich) and ionomycin (1.5 μM, Sigma-Aldrich) for 5 hours followed by cytokine staining.

### qRT-PCR

Total RNA was isolated from sorted splenic NKT and CD4 T cells from C57BL/6 mice using RNeasy Plus Mini Kit (QIAGEN) following the manufacturer’s instructions. 50 ng total RNA was used to do single-strand cDNA synthesis with the RT2 First Strand Kit (QIAGEN). Predesigned primer sets for each gene, i.e. Nox1, Nox2, Nox3, Nox4, Duox1 and Duox2 were purchased from IDT. Either ⁏-actin or GAPDH were used as internal controls. The inverse log of the delta delta-CT was then calculated.

### Statistical analysis

Data for all experiments were analyzed with Prism software (GraphPad Prism ver. 6, San Diego, CA). Unpaired Student’s t-test was used for comparison of experimental groups. Correlation coefficient (r^2^) was calculated by fitting the data to a linear regression curve. The following p values were considered statistically significant: *p < 0.05, **p < 0.01, ***p < 0.001, and ****p<0.0001.

## Results

### Peripheral NKT cells have greatly elevated ROS levels

To examine if NKT cells show different metabolism, we measured ROS levels as a parameter of cell metabolism using 2’,7’–dichlorofluorescin diacetate (DCFDA), a fluorogenic dye that is commonly used to detect hydroxyl, peroxyl and other ROS activity within the live cells (40). ROS levels in NKT cells were similar to CD4 or CD8 single positive cells in the thymus but greatly increased in the spleen and liver with a mixture of ROS high and low cells (Fig. 1A). We further characterized the NKT cell population to get a better understanding of the ROS levels and the NKT cell phenotype using CD44 and CD62L that are markers of NKT cell maturity and activation (6). We compared the levels of ROS of CD44-high and CD44-low cells as well as CD62L-hi and CD62L-low NKT cells. The results showed that CD44-high or CD62L-low splenic NKT cells had high levels of ROS, and NK1.1 expression did not correlate with ROS levels (Fig. 1B). The high levels of ROS in NKT cells did not associate with the effector phenotype or being innate type cells because neither CD44^+^ effector CD4 T cells nor NK cells showed high ROS from the spleen and liver (Fig. 1C). Thus, highly elevated ROS is a unique property of NKT cells.

**FIGURE 1.**
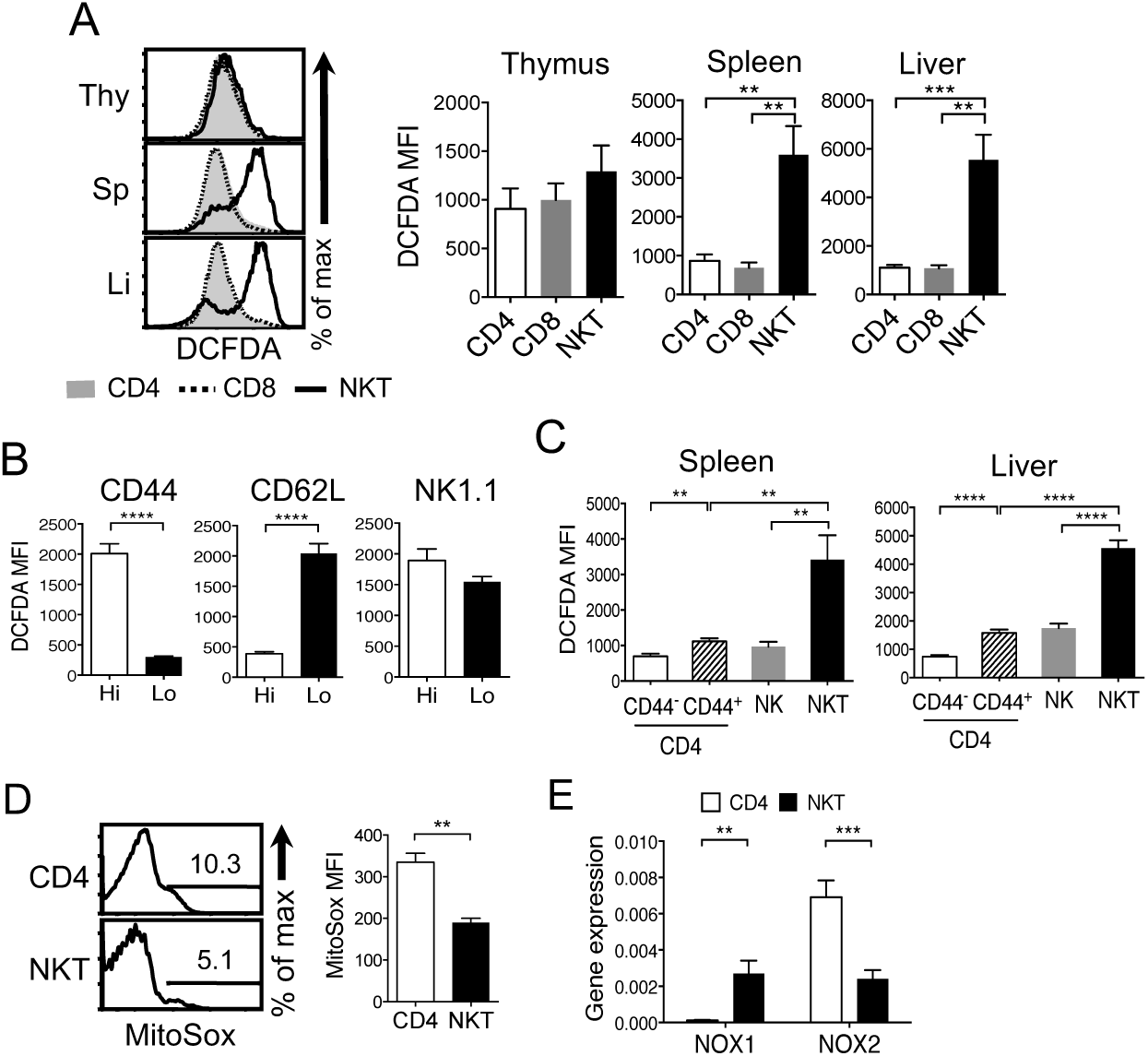
Study of ROS levels in NKT and CD4 T cells. (**A**) Freshly isolated thymocytes, splenocytes and liver lymphocytes were compared for the amount of ROS using 2’,7’-dichlorofluorescin diacetate (DCFDA) as described in Materials and Methods. NKT cells were identified by TCR-β low and CD1d-tetramer^+^. N= 8 mice per group. (**B**) Expression level of CD44, CD62L, and NK1.1 was used to compare the amount of ROS in two subsets of splenic NKT cells. N=4 mice per group. (**C**) NKT, CD44^+^ and CD44^-^ CD4 T cells, and NK cells from spleen and liver were compared for the levels of ROS. N=6 mice per group. (**D**) NKT and CD4 T cells from total splenocytes were compared for mitochondria-produced ROS using MitoSox, as described in the Methods. N=3 mice per group. (**E**) Gene expression of the members of the Nox gene family was assessed from sorted splenic NKT and CD4 T cells using qPCR. Gene expression was normalized to either β-actin or GAPDH. N=4 mice per group. All flow cytometric data were analyzed with live cells and C57BL/6 mice were used. One representative histogram is shown from at least three independent experiments. Error bars represent the mean ± SEM. **p <0.01; ***p<0.001; ****p<0.0001.

Next, we investigated the source of ROS in NKT cells. Because ROS are mainly produced by NADPH oxidases (NOX) and mitochondria (41), we first measured ROS produced by mitochondria (mtROS) using MitoSox (42). Unlike high ROS measured by DCFDA, a small fraction of NKT cells produced mtROS and both the frequency of mtROS-producing NKT cells and the amount of mtROS measured by MFI were lower than that of CD4 T cells (Fig. 1D) indicating that ROS in NKT cells are not produced by mitochondria in the steady state. To determine if ROS were produced by NOX, we compared the gene expression of members of the NOX family - Nox1, Nox2, Nox3, Nox4, dual oxidases 1 and 2 (Duox1 and Duox2) - in NKT and CD4 T cells using qPCR. CD4 T cells predominantly expressed Nox2, whereas NKT cells expressed both Nox1 and Nox2 (Fig. 1E). Nox3, Nox4 and DUOXs mRNA levels were below detection or very low (data not shown). Together, ROS in NKT cells are likely generated by NOX1 and NOX2.

### ROS-high NKT cells are susceptible to oxidative stress but not ER stress

Because excessive ROS can cause stress and damage the cells, we asked whether NKT cells are susceptible to oxidative stress. To test this, we treated the cells with hydrogen peroxide (H_2_O_2_). Appropriate concentration of H_2_O_2_ provides a permissive oxidative environment for cellular signaling which is ideal to maintain homeostasis and to adapt to the stress. However, H_2_O_2_ levels above the optimal range cause oxidative damage and aberrant cell signaling resulting in pathologies (34). We subjected total splenocytes to a range of H_2_O_2_ concentrations (10 μM to 150 μM) and measured apoptosis to compare responses. NKT cells showed increased percentages of apoptotic cell death at 30 μM, whereas a higher dose of H_2_O_2_ was required to induce cell death of CD4 T cells (Fig. 2A). The data were consistent with the reduced NKT cell frequency and low ROS in remaining live cells (Fig. 2B). If apoptosis is caused by high ROS, cell death can be prevented by reducing ROS. This was tested by treating cells with antioxidants N-acetyl-cysteine (NAC) or diphenylene-iodonium (DPI) prior to addition of H_2_O_2_. NAC is an effective free radical scavenger, while DPI inhibits the activity of NOXs and DUOXs that generate superoxide anion and hydrogen peroxide, respectively(43, 44). Both NAC and DPI pre-treatment reduced NKT cell death caused by H_2_O_2_ (Fig. 2C). However, MitoQ, a mitochondria-targeted antioxidant that inhibits mitochondrial ROS (45), failed to protect the oxidative damage caused by H_2_O_2_ (Fig. 2C), indicating that mtROS do not induce oxidative stress in NKT cells. Next, we asked if high levels of ROS in NKT cells would render sensitive responses to other cellular stress conditions. To test this, we treated NKT and CD4 T cells with different concentrations of thapsigargin that inhibits endoplasmic reticulum (ER) Ca_2_ ^+^ ATPase to induce stress (46). In contrast to H_2_O_2_ treatment, NKT cells were highly tolerant to thapsigargin, whereas CD4 T cells showed cell death at 100nM concentration (Fig. 2D). Therefore, high ROS in NKT cells does not cause sensitive responses to ER stress

**FIGURE 2.**
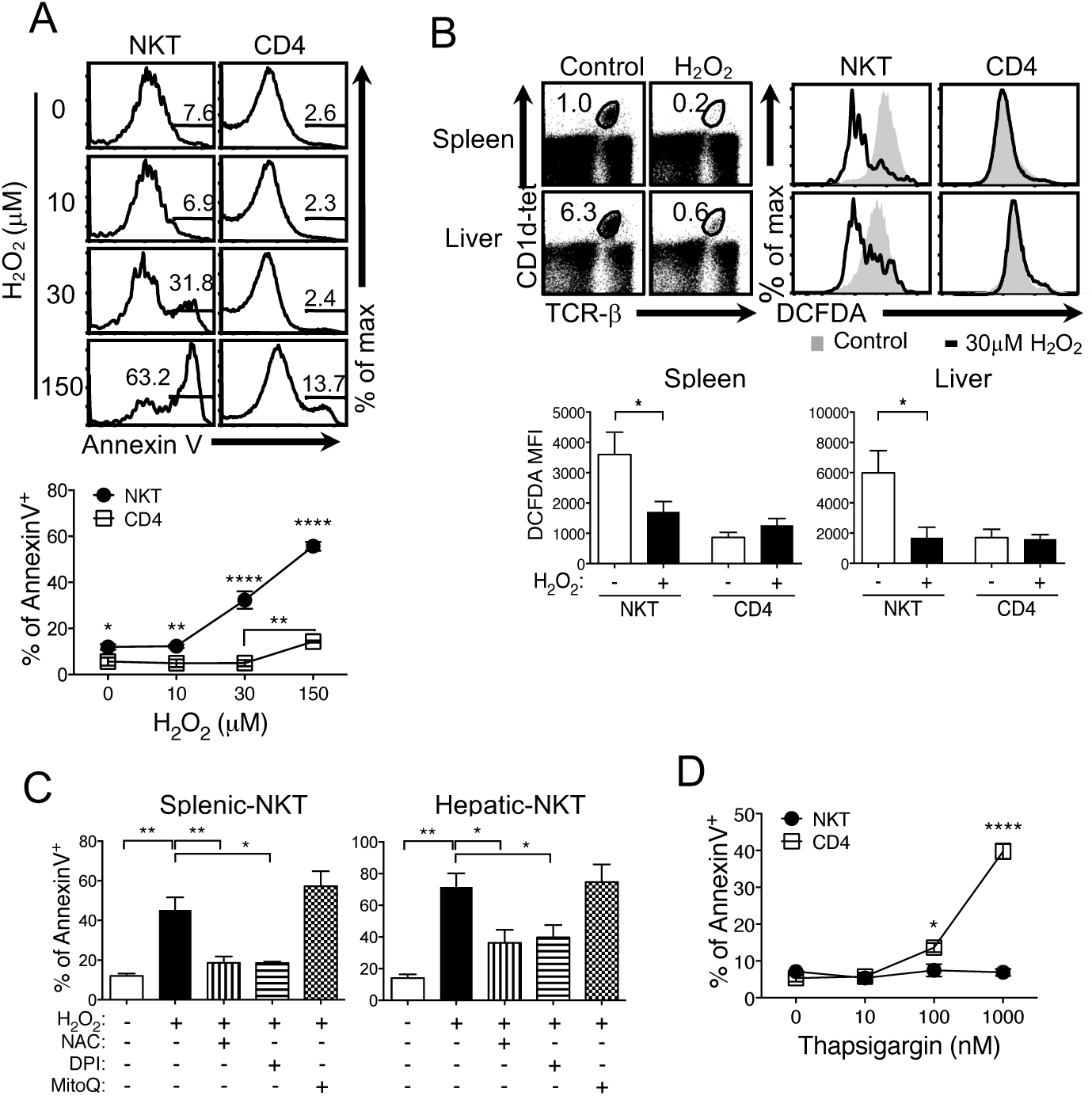
NKT cells are susceptible to oxidative stress. (A) Total splenocytes were treated with H_2_O_2_ for 30 minutes at indicated concentrations and then measured for apoptosis using Annexin V. The graph illustrates the summary of % of Annexin V^+^ cells. N=6 mice per group. (B) Total spleen and liver cells were treated without or with 30 μM H_2_O_2_ for 30 minutes and examined for the ROS levels. N=8 mice per group. (C) Total lymphocytes from spleen and liver were pre-treated with 15 mM NAC or 20 μM DPI for 30 minutes, or 100 nM MitoQ for 60 minutes and treated with 30 μM H_2_O_2_ for 30 minutes followed by the same assay as in (A). N=6 mice per group for NAC and DPI treatments and N=3 mice per group for MitoQ treatment. (D) Total splenocytes were treated with Thapsigargin for 30 minutes at indicated concentrations before measuring apoptosis. N=4 mice per group. C57BL/6 mice were used and the histograms and dot plots are representative of at least three independent experiments. Error bars represent the mean ± SEM. *p<0.05; **p <0.01; ****p<0.0001.

### The amount of ROS correlates with the function of NKT cells

As effector cells, NKT cells can produce cytokines upon short stimulation (47). Because effector functions of T cells are regulated by ROS (36, 48), we asked if ROS also control the function of NKT cells. To test this, we reduced ROS by pre-treating freshly-isolated lymphocytes with antioxidant for 30 minutes followed by stimulation with PMA and Ionomycin for 5 hours. The ROS level was decreased upon pre-treatment with 20 μM DPI in NKT cells but not in CD4 T cells from the spleens (Fig. 3A). Hepatic NKT and CD4 T cells showed the same pattern (Supplemental Fig. 1A). When cytokine expression was examined, DPI-treated NKT cells from spleen and liver had more IL-4^+^ cells but fewer IFN-γ^+^ and IL-17^+^ cells than without DPI treatment (Fig. 3B, *upper panel*; Supplemental Fig. 1B, *upper panel*). CD4 T cells did not show a measurable change with DPI treatment except an increase in IFN-γ^+^ cells (Fig. 3B, *lower panel*; Supplemental Fig. 1B, *lower panel*). Because it is possible that DPI treatment might have caused a selective loss or gain of an effector cell type, we sorted ROS-high and ROS-low NKT cells and examined effector cell subsets. Sorted ROS-high and ROS-low NKT cells were assessed for the expression of T-bet, GATA3 and RORγt, the transcription factors responsible for generation of NKT1, NKT2, and NKT17 cells, respectively. ROS-high NKT cells had more NKT1 but fewer NKT2 than ROS-low cells (Fig. 3C). Although NKT17 data did not reach a statistically significant difference, the trend agreed with the cytokine data. Together, both the cytokine data and the functional subset analyses suggest that the amount of ROS correlate with the NKT cell function.

**FIGURE 3.**
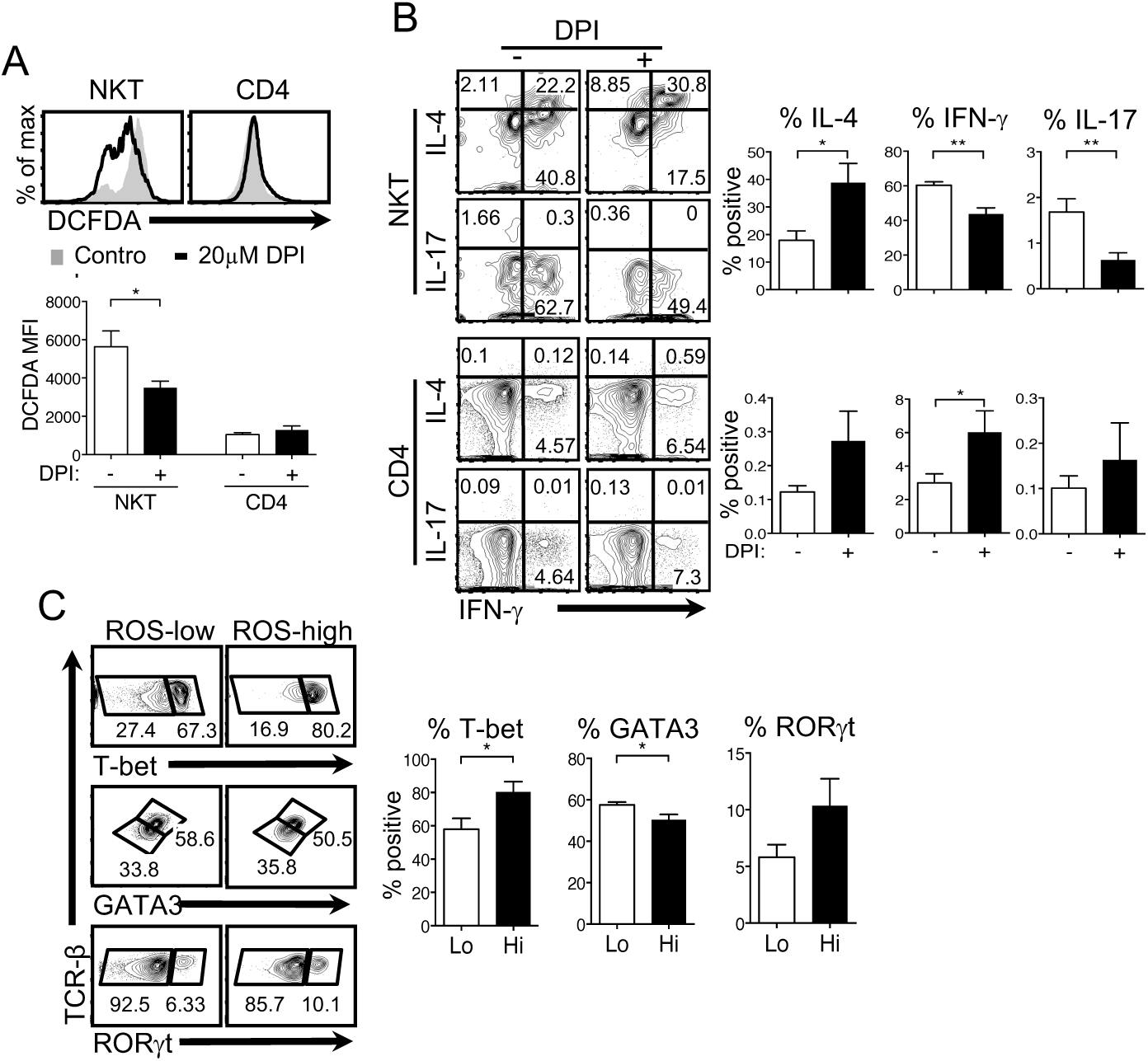
Regulation of cytokine expression by ROS. (**A**) Total splenocytes from C57BL/6 mice were treated with 20 μM DPI for 30 minutes and ROS was measured. N= 5 mice per group. (**B**) The cells pretreated with 20 μM DPI as described in (A) were washed and then stimulated with PMA (50 ng/ml) and Ionomycin (1.5 μM) for 5 hours to measure cytokine expression by NKT and CD4 T cells. N=6 mice per group. (**C**) ROS-low and ROS-high NKT cells were sorted from C57BL/6 mice spleens and analyzed for the expression of indicated transcription factors. N=4 mice per group. Each flow cytometry histogram and dot plot shows one representative experiment. Error bars represent the mean ± SEM. *p<0.05; **p <0.01.

### ROS levels of NKT cells decrease upon activation along with reduced IFN-γ+ and IL-17+ cells

It is reported that CD4 T cells increase mtROS production upon activation (36). Therefore, we asked whether NKT cells also produce mtROS upon activation further increasing total ROS. To answer this, NKT cells from Vα14 transgenic mice were used to obtain a sufficient number of NKT cells for the assays. Vα14 NKT cells had ROS levels comparable to NKT cells from C57BL/6 mice (Supplemental Fig. 2A). Splenic Vα14 NKT and CD4 T cells from C57BL/6 mice were sorted and stimulated with α-Galcer or anti-CD3 plus anti-CD28, respectively. We then compared the amounts of total ROS and mtROS before and after 3 days of stimulation. As expected, both total and mtROS increased upon activation at day 3 in CD4 T cells. In contrast, in NKT cells, total ROS levels were reduced despite a comparable increase of mtROS to CD4 T cells (Fig. 4A). Increase of CD69 expression indicates that both cell populations were activated (Fig. 4A). NKT cells showed similar response when stimulated with α-Galcer or with anti-CD3 and anti-CD28 antibodies (data not shown). Moreover, we found that CD62L-low and -high NKT cells exhibit decreased and increased ROS, respectively, after stimulation (Supplemental Fig. 2B). CD62L-high but not CD62L-low CD4 T cells also showed an increase in ROS levels (Supplemental Fig. 2B). We showed that low ROS by DPI pretreatment changed cytokine expression (Fig. 3B). Therefore, we wanted to test if cytokine expression in activated NKT cells would be similar to that of DPI pretreated NKT cells due to low ROS. As shown in Figure 4B, activated NKT cells showed less IFN-γ^+^ and IL-17^+^ but more IL-4^+^ cells than before activation. Activated CD4 T cells also showed a similar response to those treated with DPI (Fig. 4C).

**FIGURE 4.**
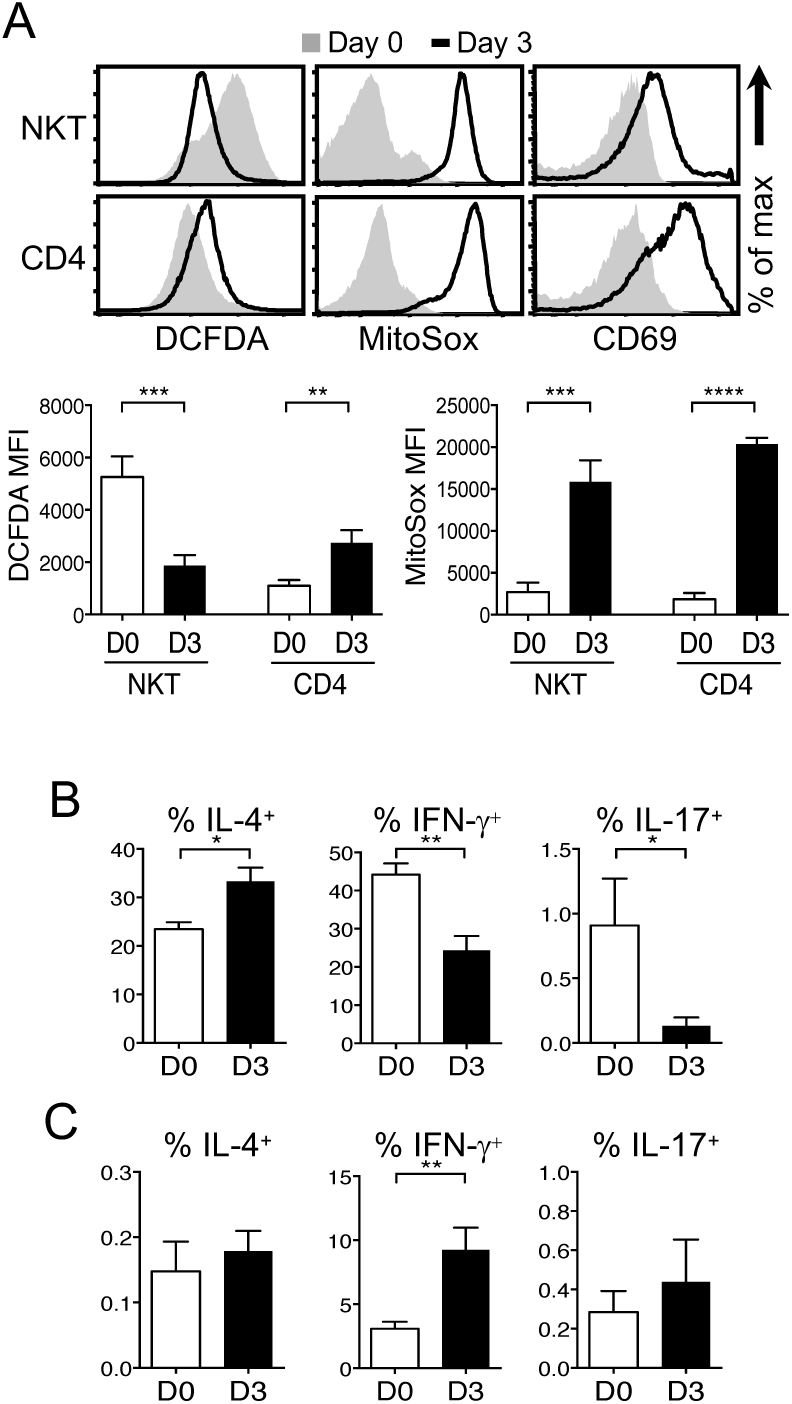
ROS levels change after activation. (**A**) NKT and CD4 T cells were sorted from Vα14 and C57BL/6 mice spleens, respectively, followed by stimulation as described in the Materials and methods for 3 days. Cells at day 0 and 3 were analyzed to measure total ROS (DCFDA), mtROS (MitoSox) and CD69 expression. N=6 mice per group. (**B** and **C**) The same cells in (A) were re-stimulated with PMA and Ionomycin for 5 hours to measure cytokine expression by NKT (B) and CD4 T cells (C). N=5 mice per group. Histograms are representative of at least six independent experiments. Error bars represent the mean ± SEM. *p<0.05; **p <0.01; ***p<0.001; ****p<0.0001.

### ROS levels correlate with expression of PLZF

It is known that NKT cells but not CD4 T cells express PLZF so we looked into a possible role of PLZF in increased ROS levels of NKT cells. To answer this, we examined γδ T cells that are known to express PLZF (49, 50). We found that splenic and hepatic γδ T cells had low and high ROS respectively (Fig 5A). To further investigate the role of PLZF in ROS, we compared the amounts of ROS with PLZF expression levels using CD4, NKT and γδ T cells from C57BL/6 mice. As shown in Figure 5B, we observed a positive correlation between PLZF and ROS levels. Cells expressing higher levels of PLZF such as NKT cells had more ROS than those with less or no PLZF, strongly suggesting that the level of ROS is controlled by PLZF. To further examine the role of PLZF in ROS levels in mice without a genetic modification, we examined NKT cells in visceral adipose tissues (VAT). A study reported that these NKT cells do not express PLZF and are different from those in other tissues in their development, phenotype and function (23, 51). We found that NKT cells from VAT (VAT-NKT) had lower levels of ROS than those from spleen or liver (Fig. 5C). Lastly, we compared PLZF expression in sorted ROS-high and ROS-low NKT cells from C57BL/6 that were used in Figure 3C. ROS-high NKT cells expressed higher PLZF than ROS-low cells (Fig. 5D), further indicating a role of PLZF in the regulation of ROS levels.

**FIGURE 5.**
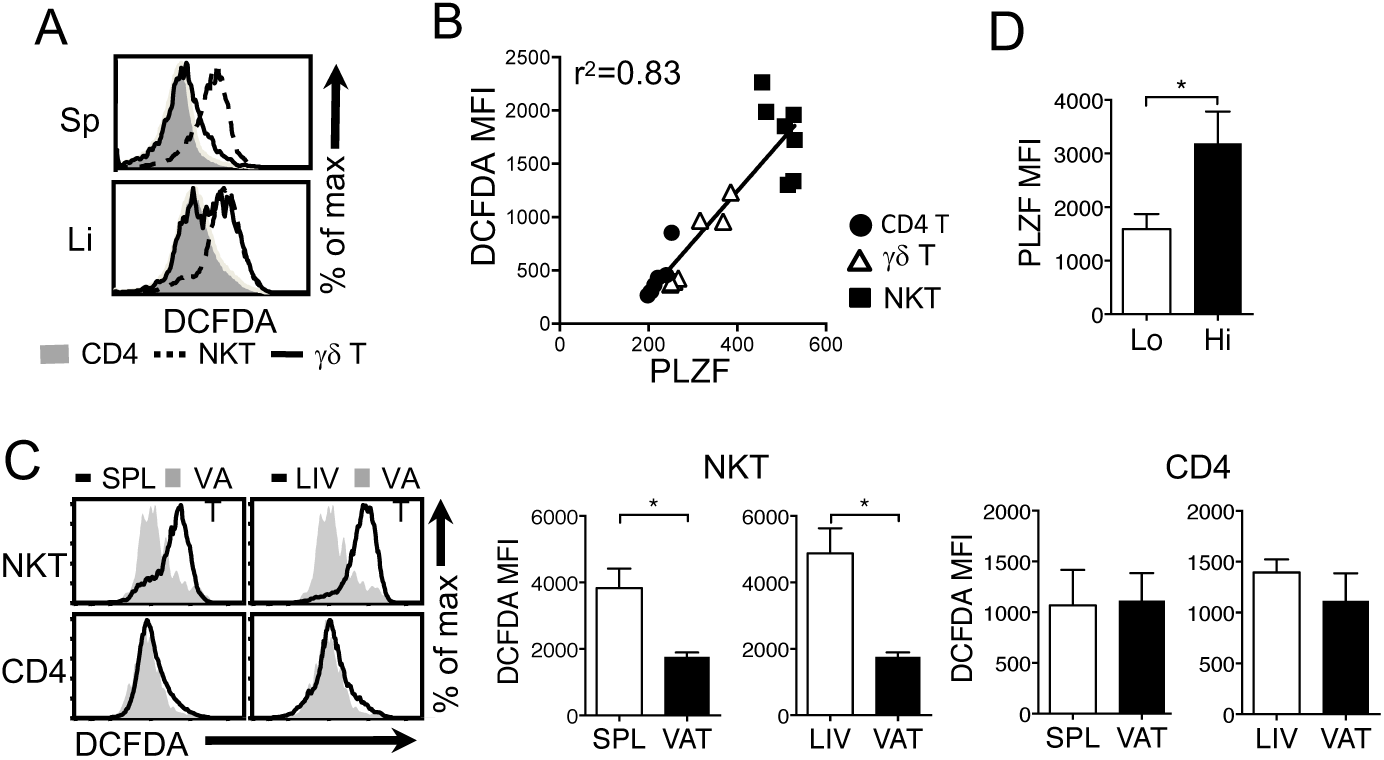
ROS levels correlate with PLZF expression. (**A**) Total lymphocytes from spleen and liver of C57BL/6 mice were used to measure ROS levels of CD4 T, NKT and γδ T cells. (**B**) The graph showing the correlation between levels of ROS and PLZF of CD4 T, NKT and γδ T cells. N=4 (spleen) or N=3 (liver) mice per group. Splenic and hepatic samples were not distinguished in the graph. (**C**) Total lymphocytes from the spleen or visceral adipose tissues (VAT) from C57BL/6 mice were used to measure ROS levels of NKT vs. CD4 T cells. N=3 mice per group. (**D**) ROS-low and ROS-high cells used in Fig 3C were examined to measure PLZF levels. N=4 mice per group. Histograms are representative of at least three independent experiments. Error bars represent the mean ± SEM. *p<0.05.

### PLZF regulates ROS levels

If PLZF controls the level of ROS, it is expected that NKT cells with reduced PLZF should have lower ROS than NKT cells with normal amounts of ROS. To test this, we examined NKT cells from PLZF haplodeficient (PLZF^+/-^) mice which express less PLZF. Total deficiency of PLZF is not informative because PLZF is required for NKT cell development (5, 6). The ROS levels were greatly decreased in NKT but not CD4 T cells from PLZF^+/-^ mice compared to the wild type (WT) mice (Fig. 6A; Supplemental Fig. 3A). Consequently, PLZF^+/-^ NKT cells were more tolerant to oxidative stress than WT NKT cells (Fig. 6B; Supplemental Fig. 3B). Next, we asked if PLZF expression is sufficient to elevate ROS levels. To answer this, we examined CD4 T cells from PLZF transgenic (PLZF^Tg^) mice under the premise that expression of PLZF in CD4 T cells that normally do not express PLZF would increase ROS levels and exhibit a sensitive response to oxidative stress similar to NKT cells. Indeed, ROS levels were greatly increased in CD4 T cells from PLZF^Tg^ mice compared to WT CD4 T cells, from both spleen and liver (Fig. 6C; Supplemental Fig. 3C) and became sensitive to a low concentration of H_2_O_2_ treatment (Fig. 6D; Supplemental Fig. 3D). NKT cells were comparable between WT and PLZF^Tg^ mice (Fig. 6C; Supplemental Fig. 3C). The induction of ROS in PLZF^Tg^ CD4 T cells was not due to an artifact associated with the line of transgenic mouse because NKT cells from an independently generated PLZF transgenic mice showed the same response (Supplemental Fig. 3E). Similar to activated NKT cells, activation of PLZF^Tg^ CD4 T cells resulted in the reduction of ROS (Fig. 6E; Supplemental Fig. 3F) and decreased IFN-γ and IL-17 expressing cells but not IL-4^+^ cells (Fig. 6F). Taken all together, PLZF regulates ROS levels, which in turn controls inflammatory function of NKT cells.

**FIGURE 6.**
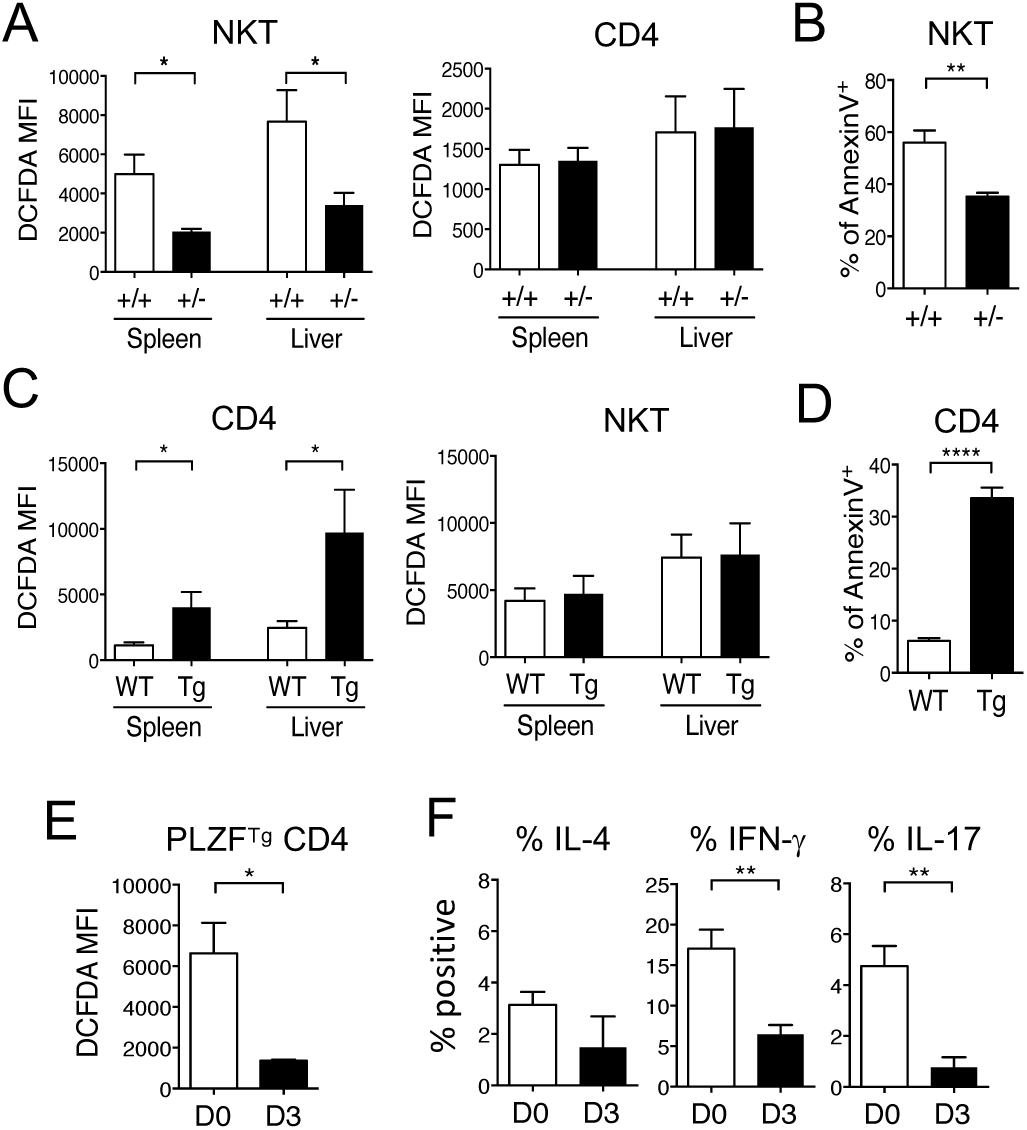
PLZF regulates ROS levels (**A**) NKT and CD4 T cells from spleen and liver of PLZF^+/-^ mice and their litter mates (PLZF^+/+^) were examined for ROS. N=5 mice per group. (**B**) Splenic NKT cells from PLZF^+/-^ and PLZF^+/+^ were measured for apoptosis after 30 μM H_2_O_2_ treatment. N=5 mice per group. (**C**) CD4 T cells from spleen and liver of PLZF^Tg^ (Tg) mice and their litter mates (WT) were compared for ROS. N=5 mice per group. (**D**) Splenic CD4 T cells from Tg and WT mice were measured for apoptosis as in (B). N=5 mice per group. (**E** and **F**) Enriched splenic PLZF^Tg^ CD4 T cells were stimulated for 3 days as described in the Materials and Methods and then measured for ROS (E) and cytokine expression (F). N=4 mice per group. Error bars represent the mean ± SEM. *p<0.05; **p <0.01; ****p<0.0001.

## Discussion

During thymic development, NKT and CD4 T cells showed little difference in ROS levels but, in the periphery, ROS were greatly elevated in NKT cells while CD4 T cells maintained the similar levels. It is not clear what causes the increase of ROS in the peripheral NKT cells, but it does not seem to be due to TCR-mediated signaling. It is reported that NKT cells receive very weak, if any, TCR stimulation in the steady-state measured by Nur77 reporter expression (51). In addition, we showed that αGalCer stimulation reduces ROS. Therefore, it is unlikely that NKT cells are continuously undergoing TCR-mediated activation leading to the increase of ROS. Although TCR stimulation of T cells induces ROS production in the mitochondria (36), our data showed that only a small fraction of freshly isolated NKT cells produced mtROS indicating that ROS were from a different source. The primary source of ROS in NKT cells seems to be NADPH oxidases and NKT cells may have a higher capacity to produce ROS using the NOX system. NK cells, a relative of NKT cells, do not have high ROS, which argues against the fact that the innate nature of NKT cells is responsible for elevated ROS. Rather, it is a unique feature of NKT cells.

Although NKT cells are effector cells, the high level of ROS is not simply due to the effector phenotype of the cells because CD44-high CD4 T effector cells did not have as high ROS as NKT cells. Instead, it seems that ROS levels increase as NKT cells mature. It is not clear why NKT cells make more ROS in the steady state because the consequence of high ROS is likely detrimental to NKT cell survival. A slight increase of ROS in a local environment may result in a selective loss of NKT cells but sparing CD4 and CD8 T cells to prevent non-specific NKT cell-driven immune responses as bystanders. To prevent spontaneous cell death *in vivo*, it is possible that NKT cells have a built-in mechanism to cope with high ROS. The response of NKT cells to ER stress was not altered implying that high ROS does not make NKT cells sensitive to other cellular stress. It is also interesting that NKT cells are more tolerant to ER stress than CD4 T cells suggesting that intracellular environment of the two cell types are different. Upon activation via TCR signaling, in contrast to CD4 T cells, NKT cells down-regulate total amounts of ROS although mtROS levels are upregulated in NKT cells suggesting different uses of ROS during activation. Perhaps ROS produced by the NOX system in the steady state of NKT cells is reduced and switched to mitochondria production of ROS. Although underlying mechanisms for this switch need to be investigated, proper control of ROS in NKT cells is important for their survival.

Reducing ROS by either TCR-mediated activation or antioxidant treatment decreased NKT cells expressing IFN-γ or IL-17 but increased IL-4^+^ cells. The role of ROS in NKT cell function is further strengthened by the results that more inflammatory NKT1 and NKT17 cells were found among ROS-high cells, whereas NKT2 cells are more abundant in ROS-low cells. It is possible that the expression of T-bet, GATA3 and RORγt is regulated by ROS-mediated signaling. Alternatively, the functional difference may reflect the heterogeneity of glycolytic potential of NKT cells. It is known that both Th1 and Th17 are highly glycolytic, which induces ROS generation (52). Th17 cells had higher expression levels of glycolytic genes and levels of glycolytic intermediates. Th1 cells were also dependent on glycolysis, but appeared to regulate this pathway post transcriptionally without accumulation of glycolytic intermediates (53). NKT cells show differential glucose uptake during maturation and effector cell generation in the thymus but ROS levels of thymic NKT cells are similar at different stages. Although we cannot rule out the possibility that glycolysis contributes to the effector cell differentiation in the thymus, it seems that the maintenance of each effector NKT subsets in the periphery requires different cell metabolism which is reflected by the ROS levels.

The current study provided several evidence demonstrating PLZF as a regulator of ROS and that the level of ROS correlates with the inflammatory and regulatory function of NKT cells. PLZF might act by regulating the expression of genes that are involved in antioxidant pathways or by directly regulating the metabolic pathways. PLZF has shown to regulate glucose homeostasis via gluconeogenesis and glucose output in the liver (10). In addition, PLZF affects lipid metabolism by exerting anti-adipogenic activity (54). It should be noted that although PLZF is a critical regulator of ROS in peripheral NKT cells, we did not observe a difference of ROS in thymic NKT and CD4 T cells, and PLZF^Tg^ CD4 T cells did not increase ROS levels in the thymus. Therefore, PLZF does not appear to control ROS levels in thymocytes. NKT cells reside in the liver and VAT, two organs that play a critical role in the development of meta-inflammation. Interestingly, NKT cells in VAT do not express PLZF (55). We showed in this study that NKT cells in the liver and VAT are ROS-high and ROS-low, respectively, which correlates with the inflammatory and regulatory function of NKT cells and the expression of PLZF.

In depth studies of mechanisms for PLZF-mediated ROS regulation and NKT cell metabolism are warranted to have a better understanding of NKT cell-mediated inflammatory and regulatory immune responses under a different local environment in vivo. However, these studies require an appropriate tool that allows deleting PLZF after NKT cell development in a tissue type specific manner. In addition, it is challenging to explore the role of ROS in NKT function in vivo. Use of pro-oxidants is toxic and administration of antioxidants would affect multiple cell types complicating the interpretation of the results. It should be noted that although clinical studies showed the beneficial effects of antioxidants to prevent and treat liver disease including chronic hepatitis C virus (56), alcoholic hepatitis or cirrhosis (57) and non-alcoholic fatty liver disease (NAFLD) (58), the therapeutic effect of antioxidants on NKT cells has not been demonstrated. Without long-term longitudinal studies, it is difficult to assess the clinical effect of antioxidant. In sum, we demonstrated for the first time that ROS controls NKT cell function and that PLZF is a critical regulator of ROS.

## Acknowledgements

We thank Dr. Phil King for his critical reading of the manuscript and insightful comments. The National Institutes of Health Tetramer Facility provided CD1d tetramers.

### Competing financial interests

The authors declare no competing financial interests.

